# CircCNNs, a convolutional neural network framework to better understand the biogenesis of exonic circRNAs

**DOI:** 10.1101/2024.07.12.603307

**Authors:** Chao Wang, Chun Liang

## Abstract

Circular RNAs (circRNAs) as biomarkers for cancer detection have been extensively explored, however, the biogenesis mechanism is still elusive. In contrast to linear splicing (LS) involved in linear transcript formation, the so-called back splicing (BS) process has been proposed to explain circRNA formation. To investigate the potential mechanism of BS via the machine learning approach, we curated a high-quality BS and LS exon pairs dataset with evidence-based stringent filtering. Two convolutional neural networks (CNN) base models with different structures for processing splicing junction sequences including motif extraction were created and compared after extensive hyperparameter tuning. In contrast to the previous study, we are able to identify motifs corresponding to well-established BS-associated genes such as MBNL1, QKI, and ESPR2. Importantly, despite prevalent high false positive rates in existing circRNA detection pipelines and databases, our base models demonstrated a notable high specificity (greater than 90%). To further improve the model performance, a novo fast numerical method was proposed and implemented to calculate the reverse complementary matches (RCMs) crossing two flanking regions and within each flanking region of exon pairs. Our CircCNNs framework that incorporated RCM information into the optimal base models further reduced the false positive rates leading to 88% prediction accuracy.

## Introduction

### Back splicing (BS) in the formation of exonic circRNAs

Different from linear RNAs containing both 5′ caps and 3′ tails, circular RNAs (circRNAs) form covalently closed loops with neither 5′–3′ polarities nor polyadenylated tails^1^. The molecular machinery for circRNA biogenesis is not well understood yet. circRNAs that are only derived from exons are mostly found in the cytoplasm, while circRNAs containing introns are mainly located in the nucleus^2^. Unlike canonical/linear splicing (LS) in the formation of linear RNAs, where the 3’ splice site of one exon is linearly ligated onto the 5’ splice site of another downstream exon, during the formation of circRNAs, the 3’ splice site of one exon is ligated onto its own 5’ splice site or the 5’ splice site of another upstream exon (see the diagrams in **Fig. S1a**)^3^. It has been reported that most circRNAs identified in human cells consist of multiple (at least two) exons^4^, which will be the focus of this study.

### Identification of circRNAs from RNA-Seq data

Due to low expression levels, circRNAs were frequently considered as mis-splicing by-products or splicing noises^1,2^. With the development of specialized library preparation protocols and tailored bioinformatics analyses, widely expressed circRNAs across species from archaea to humans have been discovered recently^1,5^. circRNA identification from RNA-Seq data essentially relies on the detection of BS reads where RNA-seq read pairs that are topologically inconsistent with the reference genomes are extracted for circRNA identification. Additional efforts based on the properties of circRNAs, including non-polyadenylation and RNase R-resistance, have also been utilized to increase the accuracy of circRNA detection^6^. Despite these efforts, apparent BS reads can also be generated from other processes including reverse transcriptase template switching during cDNA synthesis, tandem duplication of exons of the same gene, as well as trans-splicing, a process in which exons from different RNA transcripts are joined end to end^7^. Furthermore, the false positives arising from sequencing errors, alignment errors, and biased identification of circRNAs by different bioinformatics tools also make circRNA detection from RNA-seq data a very challenging task^6^.

### Machine learning applications in circRNA studies

Machine learning as a technique to learn patterns from data has been extensively applied to genomic studies in recent years. For example, to reduce the false positive rates in RNA-seq circRNA identification, CIRCPlus2 applied Gradient Boosting Decision Tree (GBDT) trained with the features extracted from read alignments to classify the candidate circRNAs detected by different tools as either true or false positives^8^. This machine learning classifier can be used as a filtering step to reduce the potential false-positive circRNAs reported by different bioinformatic pipelines.

Machine learning has also been used to classify circRNAs from other long non-coding RNAs (lncRNAs). For example, to classify circRNA transcripts from other lncRNA transcripts, support vector classifier (SVC), random forest (RF), and multiple kernel learning (MKL) were utilized in PredcircRNA^9^, and a hierarchical extreme learning machine (H-ELM) was also developed to enhance the performance of such classification^10^. A deep learning method circDeep that integrated a reverse complementary match (RCM) descriptor, an asymmetric convolutional neural network combined with bidirectional long short-term memory descriptor, and a conservation descriptor to extract high-level features was developed to distinguish between circRNAs and other lncRNAs^11^. Compared with PredcircRNA and H-ELM, circDeep increased the prediction accuracy by over 12%^11^. Nevertheless, the extraction of RCM features in circDeep is highly computationally expensive and time-consuming^12^. The rapid progress in circRNA classification against other lncRNAs using machine learning heavily depends on the accumulation of well-curated datasets^13,14^.

To date, the biogenesis mechanism of circRNAs remains poorly understood^15^. Due to the lack of a well-curated negative dataset and the susceptibility to high false positives in existing circRNA databases, insights into circRNA biogenesis through machine-learning approaches have been confined to a few recent studies. One of the earliest machine learning models attempted to elucidate circRNA biogenesis is Predicircrnatool^16^. As a support vector machine (SVM)-based model, it utilizes the conformational and thermodynamic properties of the dinucleotides in the flanking introns both upstream and downstream of the splicing sites as features. It classifies BS and LS exons (*i.e.*, constitutive exons in this case) obtained from the HEXEvent database^16,17^. Despite high model performance (*i.e.*, sensitivity of 0.907, specificity of 0.910, accuracy of 0.909, and Matthews’s correlation coefficient of 0.797) on the testing set, the negative data from HEXEvent only consisted of single exons. Given that the majority of circRNAs consist of at least 2 exons (*e.g.*, only 6.4% of 631,985 circRNAs in our study are associated with single exons), this model struggles to generalize well to situations involving multiple exons. Recently, a contextual regression network was explored to predict the potential of a randomly selected locus in the human genome to generate circRNAs^18^. This study used 6 features in the upstream and downstream region of the selected locus, including CpG islands, enhancer regions, RNA-binding protein (RBP) binding sites, simple repeats, A-to-I RNA editing sites, and DNA self-chains (alignment with itself) for model training. Despite a low prediction accuracy (*i.e.*, less than 73%), the study found that, among the examined 21,427 out of 55,689 circRNAs, the biogenesis of 43% was associated with RNA editing domain, 46% with flanking RBP binding sites, 6% with simple repeats, 3% with CpG island, and 2% with DNA self chains^18^. Apart from the suboptimal model performance, another limitation of this study was the random selection of negative training data, potentially not representative of the genuine LS sites. In another recent study where DeepCircCode was developed^19^, circRNA biogenesis was investigated by examining motifs extracted from the first convolutional layer of a trained convolutional neural network (CNN). This model distinguished circRNA (BS) exon pairs from corresponding negative (LS) exon pairs coexisting in the same transcripts but not included in the positive data based on circRNADb^20^ and circBase^21^. Admittedly, this strategy could enhance confidence in the negative dataset; however, the negative instance constructed in this manner is subject to significant influence from the circRNA databases used for constructing the positive dataset. Notably, circRNAs identified in circRNADb and circBase together represent only a small portion of the circRNAs available in existing circRNA databases (*e.g.*, 125,289 out of 2,471,730, *i.e.*, 5%, in our study). This discrepancy raises concerns about the representativeness of the negative instances, as the choice of circRNA databases used here might not capture the full diversity of circRNAs. Since the DeepCircCode model focused solely on information extraction from nearby junction sequences of splicing sites, additional complementary information such as RCMs crossing the flanking regions as well as within each flanking region of the splicing sites (namely BS/LS exon pairs) could potentially enhance model performance. Moreover, the input for the DeepCircCode model was the 200-bp sequence derived from concatenating two junction sequences (100 bp each) around the splicing sites. Consequently, the motifs learned in the middle of this 200-bp sequence might be affected by the artificial sequence concatenation.

To address the limitations of existing studies and gain a deeper understanding of exonic circRNA formations in humans through deep learning, we meticulously curated a high-quality dataset of BS and LS exon pairs. This dataset was based on by far the most comprehensive data integration of existing circRNA databases and a linear splicing database. With this high-quality dataset, we introduced CircCNNs, a convolutional neural network framework that incorporates the potential contribution of trans-factors (*i.e.*, motifs extracted from convolutional layers that can be bound by splicing-involving proteins) and cis-elements (*i.e.*, one type of RNA secondary structure data: RCM information crossing and within flanking regions of the exon pairs) to better understand circRNA formations.

We analyzed the learned sequence motifs from the first convolutional layer of CNN structures processing the splicing junction sequences. Trained on our curated dataset, our CNN models captured essential sequence motifs corresponding to well-established BS-associated genes such as MBNL1, QKI, and ESRP2. Furthermore, our model significantly reduced the false positive predictions for BS, with the highest precision of 95.41%. Finally, we found that the incorporation of RCM information contributed to a notable further reduction in false positive BS prediction. These results suggest the efficacy of our CircCNNs model framework and the high quality of our curated dataset in facilitating a better understanding of circRNA biogenesis. We deposited our dataset, model structures, RCM algorithm, as well as trained model weights in the following GitHub: https://github.com/wangc90/CircCNNs.

## Methods

### Construction of BS and LS exon pairs dataset

A recent study highlights significant disparities among the existing circRNA databases^13^. To enhance our understanding of exonic circRNA biogenesis, we manually curated a dataset of BS and LS exon pairs through the most comprehensive data integration among similar studies (see **Fig. 1)**. For the construction of BS exon pairs dataset (see **Fig. 1a**), we integrated data from the top 5 largest circRNA databases: circBase^21^ (92,375 circRNAs based on hg19), circRNADB^20^ (32,914 circRNAs based on hg19), circAltas^22^ (580,718 circRNAs based on hg38), circpedia v2^23^ (914,458 circRNAs based on hg19), and CSCD2.0^24^ (1,881,089 circRNAs based on hg38, unique to normal samples or common to both tumor and normal samples), along with 3,517 curated circRNAs from a recent study^13^. All circRNAs annotated for hg38 were converted to hg19 with pyliftover (https://pypi.org/project/pyliftover/), resulting in 580,059 circRNAs from circAltas and 1,879,701 circRNAs from CSCD2.0 respectively. After removing the duplicates based on the genomic coordinates of splicing sites, a total of 2,471,730 circRNAs remained. To identify exonic circRNAs, we filtered these circRNAs based on the exon boundaries of hg19 obtained from the UCSC table browser^25^. Specifically, circRNAs were considered exonic if the first splicing site of the circRNA was within a 5-bp distance of any exon starting positions and the second splicing site was within a 5-bp distance of any exon ending position from the same transcript (see **Fig. 1a**). The longest transcript of a gene was used to annotate each BS exon pairs. Since the majority of circRNAs involve multiple exons, we focused on circRNAs consisting of two or more exons, removing duplicated exon pairs as illustrated in **Fig. 1a**.

**Fig. 1.**
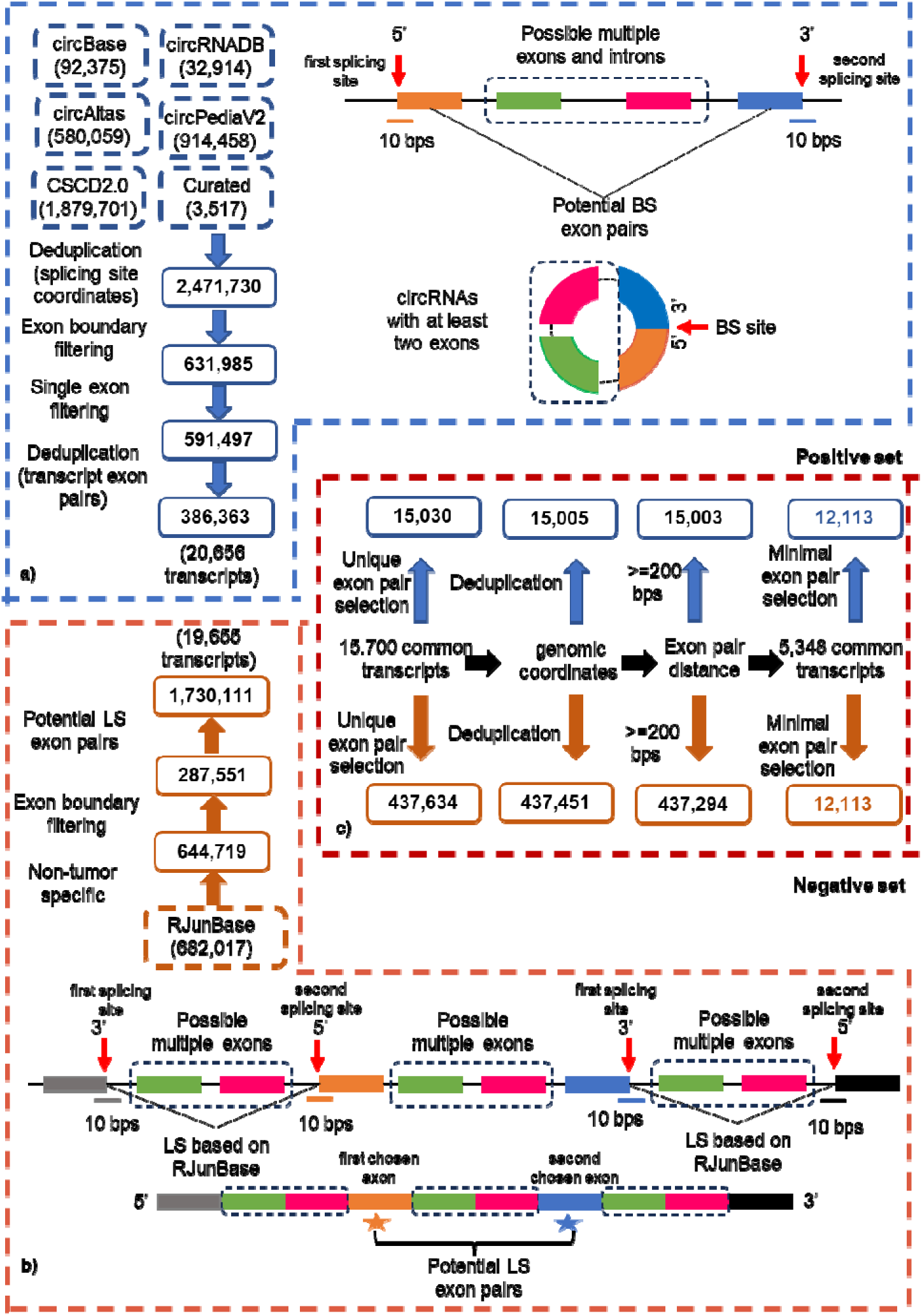
The construction of BS and LS exon pairs datasets by comprehensive data integration. a): BS exon pairs creation based on existing circRNA databases. The diagram on the top depicts the BS exon pairs and the exonic circRNAs focused by this study. b): LS exon pair creation based on RJunBase. The diagram on the bottom depicts the methods used for LS exon pairs selection based on LS evidence from RJunBase. c): two-way filtering to create the final BS and LS exon pairs dataset as positive and negative datasets, respectively.

To ensure high confidence in identifying exon pairs potentially involved only in LS (namely, LS exon pairs), we did not randomly select exon pairs not belonging to the previously obtained set of BS exon pairs, because such random selection cannot guarantee true negative data (namely LS exon pairs) if more tissue- or condition-specific RNA-Seq data indicate otherwise. In contrast, we leveraged available LS data for the human genome hg38 from the RJunBase^26^. As shown in **Fig. 1b**, we retained the LS sites that were not specific to tumors (to minimize the potential effects of genomic mutations associated with tumors on the splicing site) and converted them to hg19, resulting in 644,719 out of 682,017 reported LS sites. To confirm that these splicing sites indeed connect the 3’ end of the upstream exon to the 5’ end of the downstream exon linearly, we utilized the exon boundaries of hg19 for filtering. We retained the splicing sites and annotated the connecting exon pairs if the first splicing site was within 5 bps distance of any exon ending positions and the second splicing site was within 5 bps distance of the starting positions of any downstream exon from the same transcript (see **Fig. 1b**, note the splicing sites defined for LS differ from those for BS). The longest transcript of a gene was employed to annotate the connecting exons. Based on these connecting exon pairs, we generated potential LS exon pairs such that the 5’ of the first chosen exon is required to linearly spliced with the 3’ of any upstream exons and the 3’ of the second chosen exon is linearly spliced with the 5’ of any down-stream exons according to the RJunBase^26^ (see **Fig. 1b**). The LS exon pairs defined in this procedure are less likely to be involved in back-splicing (BS) because their splicing sites are already engaged in linear splicing (LS) and there is often a competitive relationship between BS and LS processes, where the same splicing sites may be utilized for either BS or LS^15^.

To mitigate potential bias arising from characteristics of different transcripts, we focused on comparing LS and BS exon pairs derived from the same transcript. Between the LS and BS exon pair datasets, we identified a total of 15,700 common transcripts. For each common transcript, we distinguished unique LS exon pairs, existing solely in the LS dataset, and unique BS exon pairs, found exclusively in the BS dataset (see **Fig. 1c**). Furthermore, we retained the common transcripts that have more than one exon pair of both types. Some exon pairs shared identical genomic coordinates despite being unique when considering transcript ID and exon numbers together, likely due to the overlapping transcript annotation in hg19. These redundant exon pairs were removed from both datasets. Additionally, to meet the minimum sequence length requirement for the CircCNNs, we only kept the exon pairs with at least a 200-bp separation (namely, from the 5’ of the first exon to the 3’ end of the second exon). Finally, to ensure the balanced representation of both LS and BS classes, for a given transcript, an equal number of exon pairs from each splicing type were selected (see **Fig. 1c**).

### Construction of base models

All models in this study were constructed using the PyTorch framework^27^ (version 1.13.1) with CUDA Toolkit (version 11.4) for efficient GPU acceleration. To facilitate cross-validation (CV), we employed the KFold function from sklearn^28^ (version 1.0.2) to randomly split the dataset. The Optuna (https://github.com/optuna/optuna) was used for hyperparameter searching.

Two distinct sets of models were created to incorporate the information from both the junction sequences of splicing sites and the RNA secondary structure data (*e.g.*, RCM) for classifying BS and LS exon pairs. As shown in **Fig. S1b**, we focus on RCM data crossing and within flanking regions of the exon pair. Models processing the junction sequences were termed base models, while those handling RCM data were labeled RCM models. The optimal base model and RCM model were integrated as combined models to further improve performance. As shown in **Fig. 2a**, to effectively train and evaluate these models, the BS and LS exon pairs were randomly divided into non-overlapping training and testing sets, comprising 22,000 and 2,226 exon pairs, respectively. To assess the impact of different training sizes on model performance, three distinct training sets were created from the 22,000 exon pairs: training set1 (16,000 exon pairs), training set2 (18,000 exon pairs), and training set3 (20,000 exon pairs). The remaining exon pairs were allocated to the corresponding combining sets: combining set1 (6,000 exon pairs), combining set2 (4,000 exon pairs), and combining set3 (2,000 exon pairs). All testing, training, and combining sets were balanced in the numbers of BS and LS exon pairs. Optimal hyperparameters for base models and RCM models were determined separately based on these three training sets using 3-fold CV, while optimal hyperparameters for the combined models were obtained on the combining sets.

**Fig. 2.**
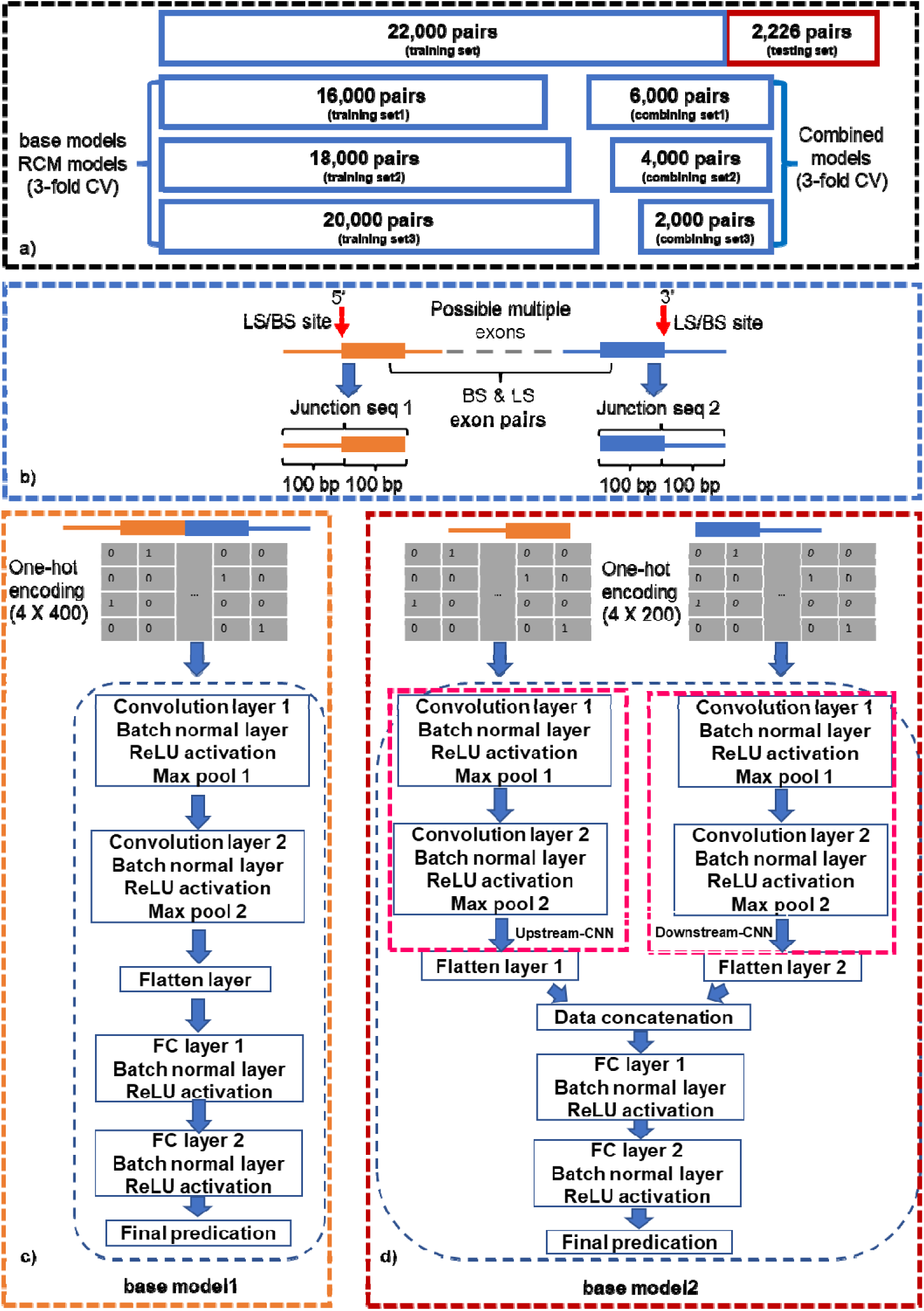
The constructions of training, combining, and testing sets, base model1 and model2. a): Separation of training, combining, and testing datasets for base, RCM, and Combined model training and final model evaluation. b): Schematic representation of the input sequences for base models. c): The architecture for base model1. d): The architecture for base model2.

Previous studies^12,19^ have underscored the importance of junction sequences around the splicing sites for distinguishing between circRNAs and lncRNAs. To select effective CNN structures capable of discerning BS from LS exon pairs based on these junction sequences, we devised two base models with distinct configurations. For base model1, mirroring the model structure of DeepCircCode^19^, a single CNN structure comprising two layers was employed. Its input sequence consists of the concatenated junction sequence around each splicing site in BS or LS exon pairs. Specifically, for each splicing site (two per exon pair), we extracted 100 bps of sequence upstream and downstream of the same splicing site (referred to as junction seq1 and junction seq2 in **Fig. 2b**). Subsequently, these two junction sequences were concatenated together and one-hot encoded to serve as the input for base model1 (see **Fig. 2c** for the model architecture). In essence, the formulation of base model1 can be expressed as follows:

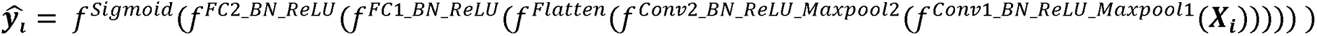

where X_i_ is the 4 X 400 one-hot encoded input matrix based on concatenated junction seq1 and seq2 shown in **Fig. 2b** for the i^th^ training example, and y--is the corresponding predicted label from base model1. For the base model2, separate CNN structures are applied to each splicing site in BS or LS exon pairs, aiming to capture the potential different sequence motifs around each site and mitigate the spurious motifs learned from the artificially concatenated sequences, a potential issue mentioned in DeepCircCode study^19^. In particular, for base model2, the input for each CNN is the one-hot encoded junction sequence from one of the splicing sites (*i.e*., junction seq1 for upstream-CNN and junction seq2 for downstream-CNN, respectively, as illustrated in **Fig. 2b** and **2d**). The outputs from these two CNNs are concatenated before the final classification, as shown in the model architecture depicted in **Fig. 2d**. Essentially, the base model2 can be expressed as follows:

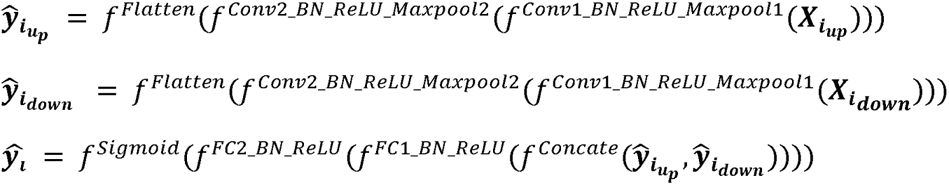

Where up stands for upstream, down represents downstream, *X_iup_* is the 4 X 200 one-hot encoded input matrix based on the junction seq1 shown in **Fig. 2b** for the i^th^ training example, and 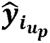 is the corresponding output from flatten layer 1 (see **Fig. 2d**). Similarly, *X_idown_* is the 4 X 200 one-hot encoded input matrix based on the junction seq2 shown in **Fig. 2b** for the same i^th^ training example, and 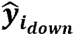 is the corresponding output from flatten layer2. 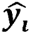 is the corresponding predicted label from the base model2. Rectified linear unit 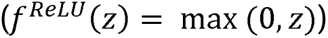 was used as the default activation function and the sigmoid function 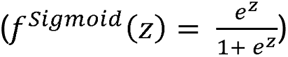 was used to make the final prediction for both base models.

The base model1 and model2 were optimized separately on each of the 3 training sets, utilizing the available choices of hyperparameters outlined in **Table S1**. The base models trained on these training sets are denoted as base1_16000, base1_18000, and base1_20000 for base model1; and base2_16000, base2_18000, and base2_20000 for base model2. The highest average 3-fold CV accuracy based on these training sets was utilized to select the corresponding optimal model hyperparameters. Given the computational constraints of exploring the entire hyperparameter space, Optuna (https://github.com/optuna/optuna), a hyperparameter optimization framework, was employed. This involved conducting 500 random trials for each model, where each trial corresponds to a different combination of hyperparameters.

### Numerical methods to detect reverse complementary matches (RCMs)

RCMs crossing the flanking intron regions of BS sites (or BS exon pairs) are believed to facilitate circRNA formation by bringing the two splicing sites from an exon pair into proximity for back splicing, typically from the 3’ end of one exon to 5’ end of an upstream exon^29^. Additionally, it has been suggested that RCMs within each flanking region of the exon pair (*i.e.*, the upstream or downstream intron of one exon pair) are involved in linear splcing^15^. Consequently, an RCM within one flanking region may compete with RCMs crossing the flanking regions, potentially influencing the likelihood of BS or LS for a given exon pair (see **Fig. S1b)**.

A previous study on the classifying between circRNAs and lncRNAs demonstrated the importance of the RCM abundance crossing the flanking regions of splicing sites^11^. To integrate the RCM model with the base models, we modified our previously established fast numerical methods in inverted repeat detection^30–32^ to determine RCM kmer pairs crossing the flanking regions as well as within each flanking region of the BS/LS exon pairs. Briefly, by converting nucleotide bases to imaginary numbers coupled with fast vector calculation, we are able to obtain the cumulative scores of kmers of any given K, thus significantly reducing the searching space for potential RCM kmer pairs. We implemented the method in a Python class and utilized the ray library (https://github.com/ray-project/ray) for parallelization.

To calculate the cumulative score for all kmers of length K in a given sequence of length L (denoted as S in **Fig. 3a**), we first convert all the nucleotide bases to a vector of real and imaginary numbers using the following mapping rule: A: 1, T: -1, C: 1j, G: -1j, N:0. This transformation is also applied to the corresponding augmented sequence (*i.e.*, “N”+sequence, denoted as AS in **Fig. 3a**) of length L+1. Then, we compute the cumulative sum vectors for both S and AS. The cumulative sum vector for sequence S represents the sum of each element up to that position, and similarly for sequence AS. Next, we calculate the cumulative score for all kmers of length k. We obtain a slice of the original cumulative sum vector starting from position K to position L, denoted as vector V1, and a slice of the same length (L-K+1) in the augmented cumulative sum vector starting from position 1 to position L-K+1, denoted as vector V2. The element-wise difference between V1 and V2 (*i.e.*, V1-V2) generates the cumulative score vector (denoted as C in **Fig. 3a**) with length L-K+1 for all kmers of length K for this given sequence S.

**Fig. 3.**
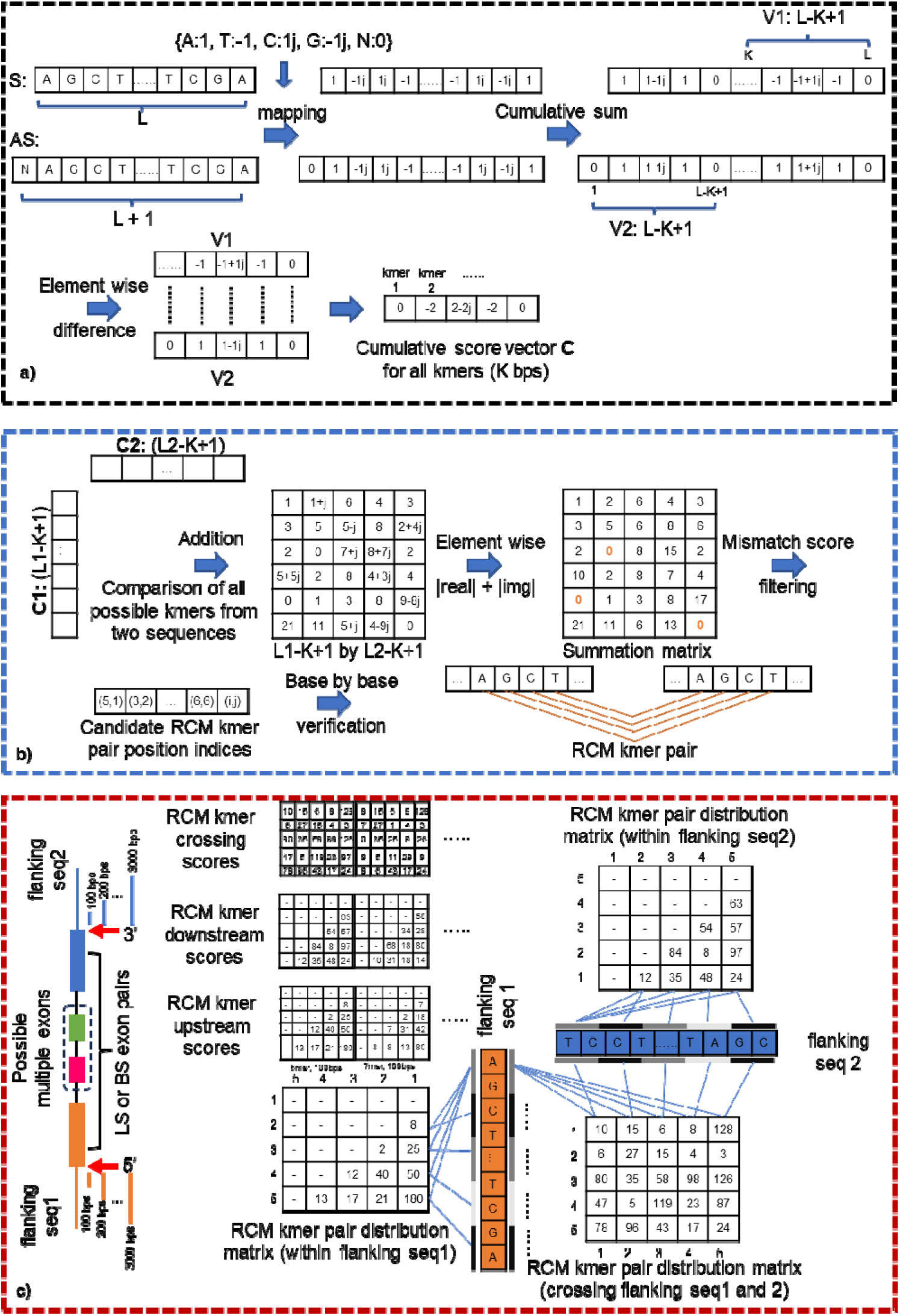
The numerical methods to determine the distribution information of RCM kme pairs crossing the flanking regions as well as in each flanking region of BS or LS exon pairs. a): Cumulative score vector creation for one input sequence. b): Determination of RCM kmer pairs based on the cumulative score vectors. c): The creation of RCM kmer crossing, upstream, and downstream scores based on the RCM kmer pair distribution matrix.

To find the RCM kmer pairs of length K crossing sequence 1 of length L1 (namely, upstream intron of the exon pair) and sequence 2 of length L2 (namely, downstream intron of the exon pair), we first obtain the cumulative score vectors for both L1 (length of L1-K+1, denoted as C1 **in Fig. 3b**) and L2 (length of L2-K+1, denoted as C2 in **Fig. 3b**) separately. These cumulative score vectors represent the potential presence of RCMs of length K at each position along the respective sequences. Next, we reshape these cumulative score vectors into column vectors of dimension (L1-K+1) by 1 for C1 and row vectors of dimension 1 by (L2-K+1) for C2, respectively. Then, we add these two vectors together by broadcasting to find the sum of all possible pairwise kmer scores in the matrix with dimension (L1-K+1) by (L2-K+1) (see **Fig. 3b**). To find candidate RCM kmer pairs crossing sequence 1 and sequence 2, we add the absolute values of the real part and imaginary part of each element in this summation matrix (see **Fig. 3b**). For each entry in this summation matrix if the sum is less than a predefined mismatch score, we define this kmer pair as a candidate RCM kmer pair crossing sequence 1 and sequence 2. We then extract the position indices of the corresponding candidate RCM kmer pairs and verify them base by base to get the final RCM kmer pairs crossing the two sequences (see the diagram in **Fig. 3b**). Similarly, to find the RCM kmer pairs within a sequence of length L, we follow the same steps with sequence 1 and sequence 2 replaced by the same sequence.

After identifying RCM kmer pairs crossing two sequences or within a sequence, we can extend these kmer pairs inwardly to find all extended subsequences that are reverse complementary to one another, constrained by some predefined overall mismatch scores. Nevertheless, a previous study indicated the absolute number of RCM kmer pairs is more informative than the longest RCM subsequences in the classification between circRNAs and lncRNAs^11^. Therefore, for each BS or LS exon pair, we obtained the RCM kmer pairs crossing the flanking intron regions as well as those within each flanking intron region of the splicing sites. For comprehensiveness, the flanking regions were defined with various window sizes ranging from 100 bps to 3000 bps (see the diagram in the left part of **Fig. 3c**). For a given window size with a certain kmer length, we determined the distribution of RCM kmer pairs crossing the flanking region and within each flanking region by defining 5 equal-distance segments along the sequence and tallying the frequency of detected RCM kmer pairs in each region represented by a 5 X 5 matrix (see **Fig. 3c**). In the previous study, to find RCM kmer pairs, the number of required comparisons is exponential as kmer length increases (*i.e.*, 4^k), making the determination of longer RCM kmer pairs computational prohibitive^11^. In contrast, our fast numerical methods enable us to handle larger kmer lengths easily. We used kmer lengths of 5, 7, 9, 11, and 13 bps without allowing any mismatches. To incorporate the distribution information of RCM kmer pairs into the model effectively, we concatenated the RCM kmer pair distribution matrices crossing the flanking regions of the splicing sites for each window size and kmer length, resulting in a 5 x (25 x # of windows size) matrix (5 different kmer lengths per window, denoted as RCM kmer crossing scores in **Fig. 3c**). Similar concatenations were performed for the RCM kmer pair distribution matrices in the upstream and downstream flanking regions of the splicing site separately (see RCM kmer upstream scores and downstream scores respectively in **Fig. 3c**). To facilitate the model training, all RCM kmer scores were log-transformed after adding 1 to avoid logarithm of 0.

### Construction of RCM models

To evaluate the relative importance of the RCM distribution within different window sizes of the splicing sites, 3 separate sets of window sizes were used. The first set includes the flanking sequences with lengths of 100, 200, 300, 400, and 500bps (denoted as small), the second set with lengths of 1,000, 1,500, 2,000, 2,500, and 3,000bps (denoted as large), and the last set with lengths defined in both small and large sets (denoted as all). To investigate the relative importance between RCM crossing and within each flanking region of the exon pairs, two different RCM models were created for each set of window sizes: (1) the first model used only the RCM kmer crossing scores as the input (denoted as RCM_crossCNN, see **Fig. 4a**) and (2) the second model used RCM kmer crossing, upstream and downstream scores together as the input (denoted as RCM_triCNN, see **Fig. 4b**). The hyperparameters for these 6 different settings (3 different sets of window size X 2 RCM models) were optimized separately on each of the 3 training sets. This optimization was done via 500 random trials using Optuna, following the same training strategy as shown in **Table S1**. The optimal hyperparameter set was chosen for the model with the highest average 3-fold CV accuracy.

**Fig. 4.**
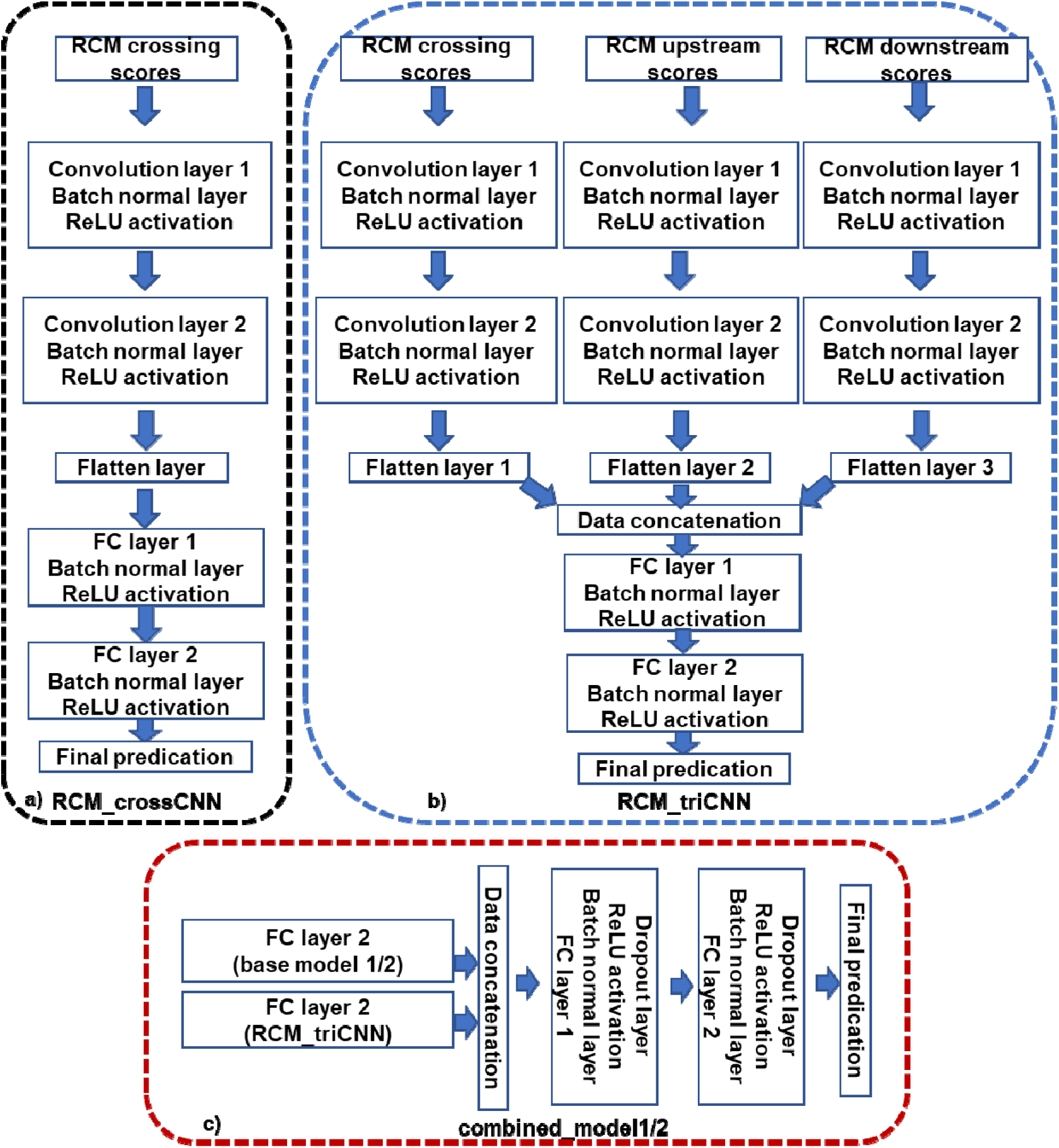
RCM models and combined models’ construction. a): Architecture for RCM_crossCNN model that only takes RCM crossing scores as the input. b): Architecture for RCM_triCNN model that takes RCM crossing, upstream and downstream scores as the input. c): Architecture to integrate the base model with the RCM_triCNN model.

### Motif extraction from retrained optimal base models

After optimizing the hyperparameters for each base model on each training set, the optimized hyperparameter set was used to retrain the models on the corresponding entire training set. Subsequently, sequence motifs potentially important for the classification task were extracted from each retrained model. This was done by applying the model weights to BS or LS exon pairs in the corresponding entire training set separately. To extract the motif features learned by each base model in the first CNN layer, we applied the motif visualization techniques used in the previous study^33^. Briefly, by using the *pytorch forward_hook*^27^ functionality, we obtained the activation values from the first convolutional layer for all the channels with the BS exon pairs in each training set as the model input. For each channel, we obtained subsequences corresponding to the maximal positive activation value with a length equal to the optimal kernel size in each model. Since we used the same-padding approach, we restricted our subsequences to the junction sequences to avoid interference from the padded regions. These subsequences were used to generate position probability matrices (PPMs), representing the proportion of 4 nucleotides in each position along the subsequences for each channel. These PPMs were then searched against the Ray2013 *Homo sapiens* RNA motif database using Tomtom (https://meme-suite.org/meme/tools/tomtom) for similarity search, utilizing the Pearson correlation coefficient. Significant matches were selected based on an E-value < 0.05 as done in the previous study^19^. The learned motifs from each retrained base model were then compared. The same motif extraction strategy was also applied to LS exon pairs to identify motif-associated genes potentially important for LS.

### Integration of retrained optimal base models with RCM models

After obtaining the optimal base model structures to process junction sequences and the optimal RCM model to process RCM information, the base and RCM models were integrated to further improve the model performance. Specifically, after retraining on each training set (see **Fig. 2a**), the model weights in the optimal base and RCM models were fixed from the convolutional layers to the second fully connected (FC) layer. These optimized model weights capture the higher-level information on junction sequences and RCMs, respectively (see **Fig. 2 and 4** for the structures of the base and RCM models). Subsequently, the outputs of the FC layer 2 from both base and RCM models are concatenated. This concatenated output is then passed through two additional FC layers and a final classification layer, as illustrated in **Fig. 4c**. These additional layers are crucial for integrating the higher-level information extracted by the previously optimized model weights. As shown in **Table S1**, the structures of these 2 additional FC layers and the corresponding dropout rates are optimized using the same training strategy. 3-fold CV with 500 random Optuna trials is conducted on the corresponding combining sets (see **Fig. 2a**) to select the optimal combining strategy for combined models that achieves the highest average CV accuracy.

### Evaluation of model performance on testing set

To ensure an unbiased evaluation of the model performance, a separate testing set (as depicted in **Fig. 2a**) that has not been involved in the model training, retraining, and model combining was utilized. To quantify the performance of retrained base models, RCM models, and combined models, a range of widely used performance metrics were employed. These metrics include ACC (prediction accuracy), specificity, recall, precision, FPR (false positive rate), FNR (false negative rate), MCC (Matthews’s correlation coefficient), and F1 score. The calculation of these metrics is based on the TP (true positives), TN (true negatives), FP (false positives), and FN (false negatives) derived from the confusion matrix on the testing set as follows: 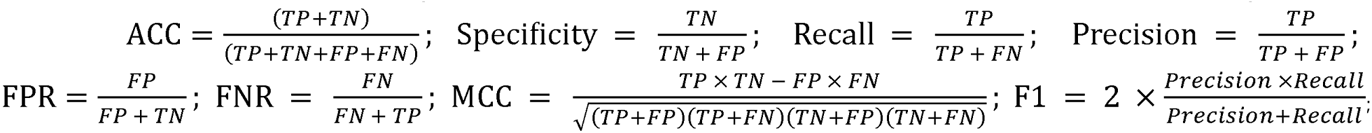 In addition, AUROC (area under the ROC curve) and AUPRC (area under the precision-recall curve) which are based on different classification thresholds were also used for performance comparison.

## Results

### Resulting BS, LS exon pairs dataset

As shown in **Fig. 1a**, after exon boundary filtering, 1,839,745 circRNAs were annotated as non-exonic circRNAs with the remaining 631,985 as exonic circRNAs. Among them, 40,467 are associated with single exons and 21 have problematic, flipped exon pairs (*i.e.*, the first exon number is larger than the second exon number). After removing those with single exons and flipped exon pairs, there are a total of 591,497 exonic circRNAs that consist of at least 2 exons. Although all of these remaining circRNAs are unique in terms of their splicing sites (*i.e.*, genomic coordinates), many of them are essentially annotated to the same exon pairs due to the slightly different splicing site annotations (*i.e.*, 1 or a few nucleotides shift) for the same circRNAs in different circRNA databases. This phenomenon was also noted by the previous study^13^. After removing these duplicated exon pairs, a total of 386,363 BS exon pairs corresponding to a total of 20,656 unique transcripts were left for further analysis. For the creation of LS exon pairs, as shown in **Fig. 1b**, after exon boundary filtering, we obtained a total of 287,551 connecting exon pairs with LS evidence based on RJunBase^26^. Based on these connecting exon pairs and according to the rationale described in the method part (also shown in the bottom diagram in **Fig. 1b**), we created a total of 1,730,111 potential LS exon pairs corresponding to a total of 19,655 unique transcripts.

We focused on the common transcripts that contain both BS and LS exon pairs and obtained 437,634 LS and 15,030 BS exon pairs, as shown in **Fig. 1c** (notice that circRNA exon pairs reduced significantly from 386,363 to 15,030). The remaining exon pairs were filtered to remove the duplicates with highly similar genomic coordinates. Consequently, 15,005 BS exon pairs and 437,451 LS exon pairs are left (see **Fig. 1c**). To satisfy the minimum sequence length requirement for the CircCNNs, we only kept the exon pairs where two exons are at least 200bps apart based on their genomic coordinates, resulting in a total of 15,003 BS and 437,294 LS exon pairs from 5,348 common transcripts. To ensure the balance of two classes, for a given transcript, we chose the same number of exon pairs from two datasets (see **Fig.1c**), resulting in 12,113 exon pairs for each splicing type, which serves as the final BS and LS exon pairs dataset for the subsequent analysis (see **Table S2** for BS, LS exon pairs dataset used in this study).

### Optimal base and RCM models

Based on the results of 500 Optuna trials for each base model on each training set (as depicted in **Fig. 2a**), the optimal base models were selected and then used for subsequence motifs extraction. As shown in **Fig. 5a**, base model2 generally exhibited slightly better performance compared to base model1 across all three training sets, with model performance improving with an increase in the number of training examples. For instance, with training set1, the optimal base model1 achieved an average 3-fold CV accuracy of 0.8596 on trial 351, while the optimal base model2 achieved a slightly higher accuracy of 0.8611 on trial 414. The optimal hyperparameters for the base model1 and 2 on each training set can be found in **Tables S3** and **S4,** respectively.

**Fig. 5.**
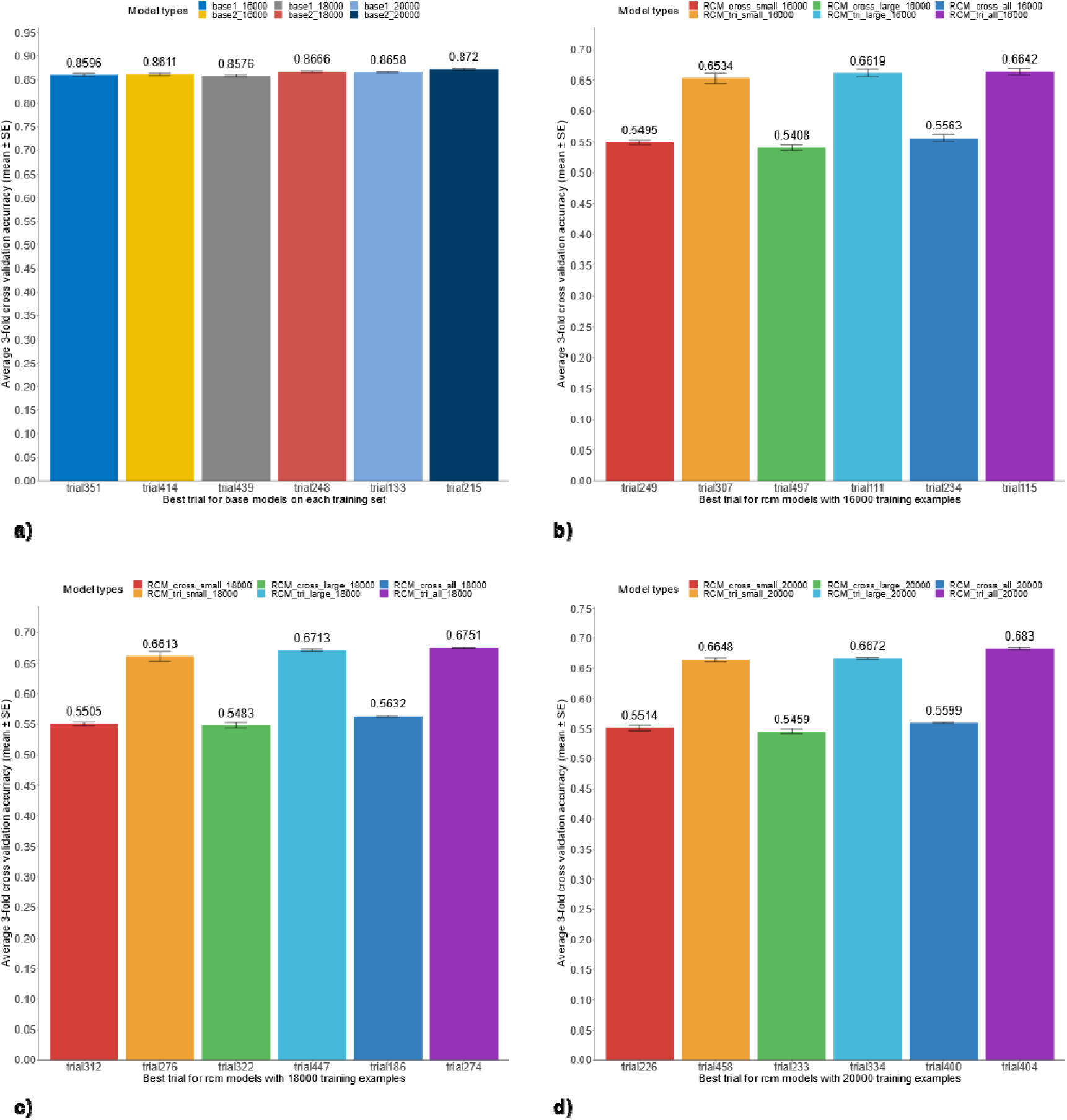
Average 3-fold CV accuracy for the optimal base and RCM models on each training set. a): average 3-fold CV accuracy for optimal base models trained on different training sets (*e.g.*, base1_16000: base model1 trained on training set1 that contains a total of 16,000 BS and LS exon pairs). b): average 3-fold CV accuracy for optimal RCM models on training set1 (RCM_cross: RCM_crossCNN model, RCM_tri: RCM_triCNN model; small: flanking sequence lengths of 100, 200, 300, 400, and 500bps were used for the determination of RCM scores; large: flanking sequence lengths of 1,000, 1,500, 2,000, 2,500, and 3,000bps were used for the determination of RCM scores; all: flanking sequence lengths of 100, 200, 300, 400, 500, 1,000, 1,500, 2,000, 2,500 and 3,000bps were used for the determination of RCM scores; RCM_cross_small_16000: RCM_crossCNN model trained on training set1 that only take RCM crossing scores derived from flanking sequence lengths of 100, 200, 300, 400, and 500bps as the input). c): The average 3-fold CV accuracy for optimal RCM models on the training set2. d): The average 3-fold CV accuracy for optimal RCM models on the training set3.

In terms of RCM models, as depicted in **Fig. 5b-d**, the RCM_triCNN model (denoted with RCM_tri_), which incorporates RCM crossing, upstream and downstream scores as model inputs (see **Fig. 3c**, **Fig. 4b**), demonstrated consistent significantly better performance compared to RCM_crossCNN model (denoted with RCM_cross_), which only considers RCM crossing scores as model inputs. For instance, the lowest average 3-fold CV accuracy achieved by the RCM_triCNN model was 0.6534 on training set 1, utilizing RCM information derived from small window sizes. In contrast, the highest average CV accuracy achieved by the RCM_crossCNN model was 0.5632 on the training set2 utilizing RCM information derived from all window sizes (see **Fig. 5b** and **5c** respectively). Given the significantly higher performance of the RCM_triCNN model across all window sizes and training sets, we then focused on the RCM_triCNN model with all window sizes in the subsequent analysis. The optimal hyperparameters for the RCM_triCNN model on each training set are provided in **Table S5.**

### Extracted motifs from retrained optimal base models

By applying the model weights of the retrained optimal base models to BS exon pairs, we obtained activation values from the first convolutional layer, which serve as sequence motif scanners for identifying sequence motifs potentially important for BS. Specifically, in the first convolutional layer of optimal base1_16000, there are 512 different channels with a size 4 X 13 (see **Table S3** for the optimal configuration of base model1 on training set1, with the optimal channel number and kernel size highlighted in bold). With the same padding strategy and a stride of 1, a total of 512 (channels) X 8,000 (BS exon pairs) X 400 (input length) activation values were obtained. Focusing on the junction sequence regions, the maximal positive activation value was identified for each training instance. For each channel and all BS exon pairs in the training set, subsequences of length 13 (since kernel length is 4 X 13 in this case) were extracted from the input sequence based on the identified maximal positive activation values. Subsequently, a position probability matrix (PPM) was obtained for each channel based on these extracted subsequences. In total, 512 PPMs were obtained for base1_16000, each with the dimension of 4 X 13 (4 nucleotides X subsequence length). Similarly, 512 PPMs were obtained for base1_18000, each with the dimension of 4 X 13; and 512 PPMs were obtained for base1_20000, each with the dimension of 4 X 17. PPMs were generated similarly for upstream- and downstream-CNNs of base model2 separately (see **Table S4** for the optimal configuration of base model2 on each training set, with the optimal channel number and kernel size highlighted in bold).

These PPMs were searched against the Ray2013 *H. Sapiens* database for motif similarity comparison^19^. Using the same approach mentioned above, we also found the sequence motifs potentially associated with LS by using the LS exon pairs as the model input. The heatmap in **Fig. 6a** summarized the presence of motif-associated genes found by different retrained optimal base models. As indicated by the columns, the PPMs extracted from all optimal retrained base models collectively matched with 45 genes from the RNA motif database with statistical significance. Each color-coded row in **Fig. 6a** represents the presence of motif-associated genes extracted from the corresponding retrained optimal base model. As shown in **Fig. 6a**, genes associated with both LS and BS were color-coded with red and labeled with “Both”. Genes exclusively associated with LS were color-coded with gold and labeled with “LS”. Genes exclusively associated with BS were color-coded with blue and labeled with “BS”. Genes identified from other retrained base models were color coded with gray and labeled with “Other”. Comparing base model1 and 2, different model structures identified different sets of genes important for LS or BS, nevertheless, some genes were recurrently identified among different model structures. For example, RBFOX1, ZC3H10, ENOX1, and MBNL1 were identified as important for both LS and BS with base model1 and model 2. RBM6 was identified as important for both LS and BS with upstream-CNN of base model2, while SNRPA was identified by downstream-CNN. Genes identified as exclusively important for either BS or LS are less consistent across different models. The extracted motifs identified as exclusively important for either BS or LS from the retrained optimal base model1 were plotted together with corresponding motif logos from the RNA motif database in **Fig. 6b** with coded color matching the legend in **Fig. 6a**.

**Fig. 6.**
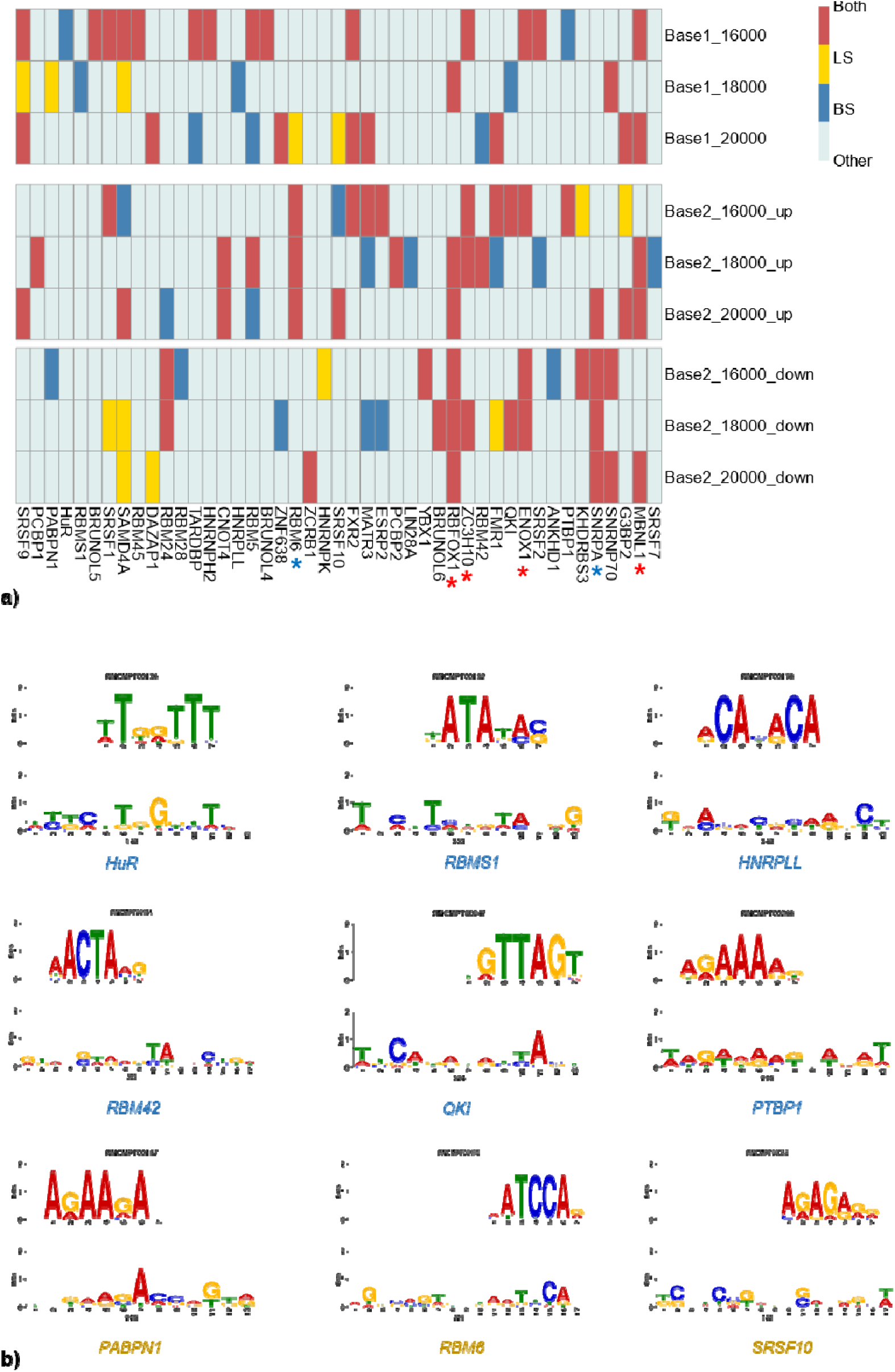
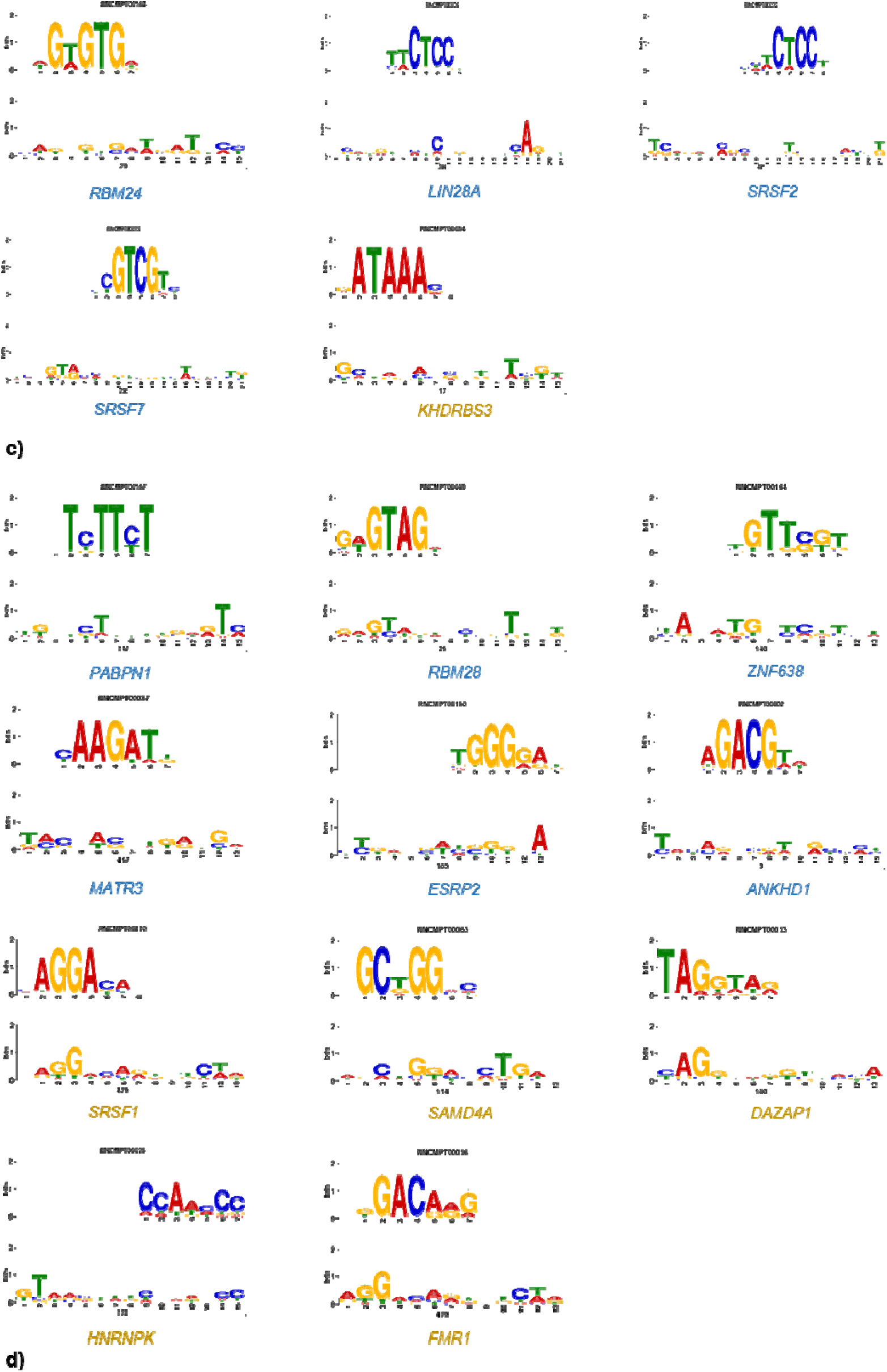
Comparisons of subsequence motifs extracted from retrained optimal base models. a): The presence of motif-associated genes extracted from retrained optimal base models stratified by whether the genes were found when both BS and LS exon pairs were used as the input or exclusively when either BS or LS exon pairs were used as the input. b): The potential genes that are exclusively associated with either BS (labeled with blue) or LS (labeled with gold) extracted from retrained optimal base model1. c) and d): The potential genes that are exclusively associated with either BS (labeled with blue) or LS (labeled with gold) extracted from upstream- and downstream-CNN of retrained optimal base model2 respectively. (The top rows in b), c) and d) represent motif logos from Ray2013 *H. Sapien* RNA motif database and the bottom rows represent the corresponding statistically similar motifs extracted from retrained optimal base models)

With upstream-CNN of base model2, genes including RBM24, LIN28A, SRSF2, and SRSF7 were identified as exclusively associated with BS, while only KHDRBS3 was identified as exclusively associated with LS. With downstream-CNN, genes including PABPN1, RBM28, ZNF638, MATR3, ESRP2, and ANKHD1 were identified as exclusively associated with BS, while genes including SRSF1, SAMD4A, DAZAP1, HNRNPK, and FMR1 were identified as exclusively associated with LS. Similarly to **Fig. 6b**, the extracted motifs identified as exclusively important for either BS or LS from the retrained optimal base model2 were plotted together with the corresponding motif from the RNA motif database. **Fig. 6c** and **6d** represent the motifs extracted from the upstream- and downstream-CNN respectively.

### Optimal combined models

Among 500 Optuna trials for each combined model on the corresponding combining set (see **Fig. 2a**), the combined models with optimal hyperparameter sets were chosen. As shown in **Fig. 7**, compared with **Fig. 5a**, the integration of the additional information extracted by the RCM_triCNN model increased the model performance of both base models on all combining sets. Specifically, the integration of the RCM_triCNN model increased the average 3-fold CV accuracy of base model1 from 0.8596 to 0.872, from 0.8576 to 0.8848, from 0.8658 to 0.877 on combining set1, 2, and 3, respectively. Similarly, the integration of the RCM_triCNN model increased the average 3-fold CV accuracy of base model2 from 0.8611 to 0.8732, from 0.8666 to 0.8855, from 0.872 to 0.878 on combining set1, 2, and 3, respectively. Like the situation for the optimal base models, the combined model with base model2 achieved a slightly higher model performance on all three combining sets. Contrary to the situation for the optimal base models, where model performance generally increased with increased training examples (see **Fig. 5a**), the highest performance for the combined models was achieved on the combining set2, which contains 4,000 LS and BS exon pairs. The corresponding optimal hyperparameters for combined models on each combining set are described in **Table S6.**

**Fig. 7.**
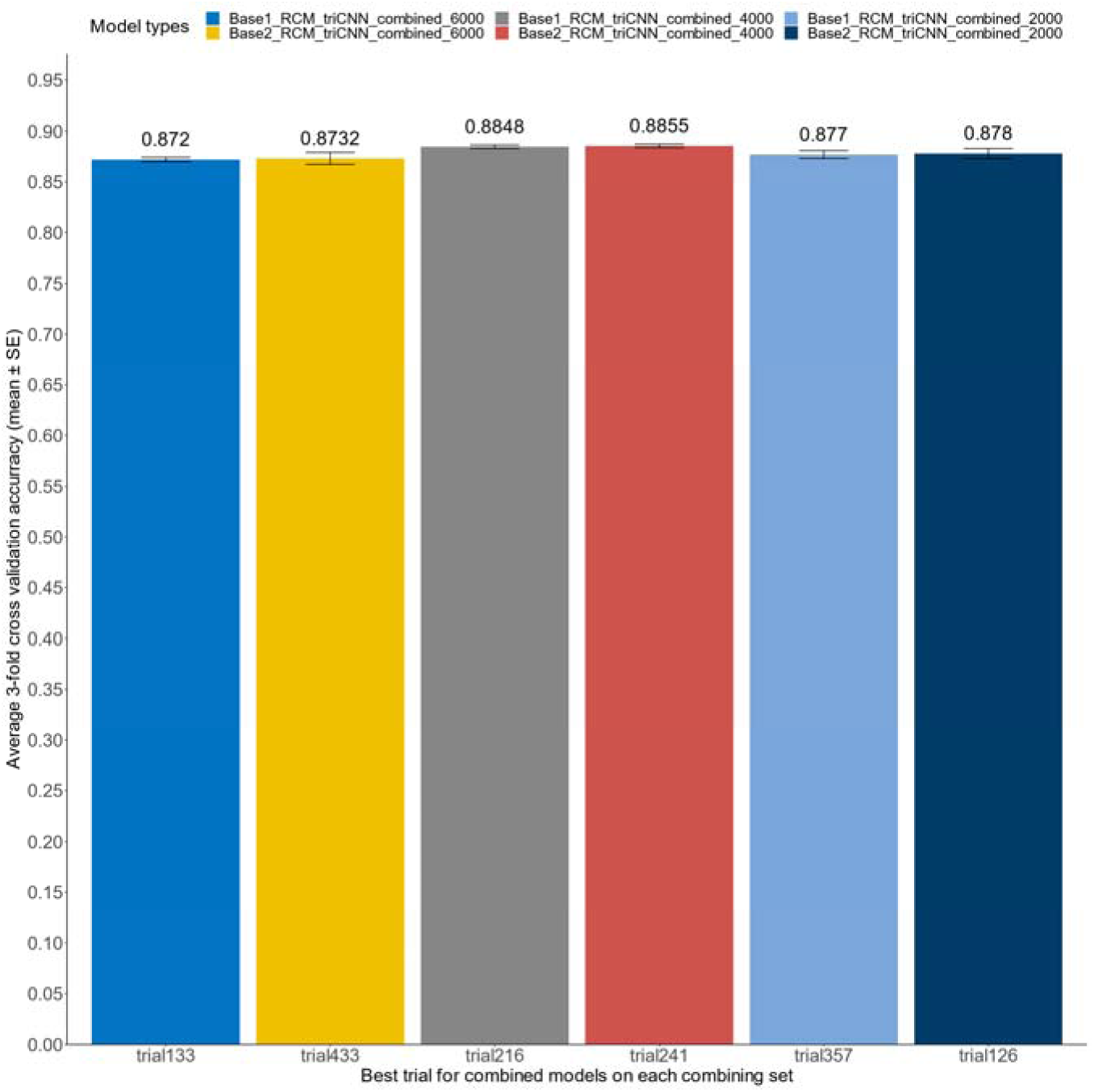
Average 3-fold CV accuracy for optimal combined models on each combining set. (Base1_RCM_triCNN_combined_6000: integration of optimal base model1 and RCM_triCNN on combining set1; Base2_RCM_triCNN_combined_6000: integration of optimal base model2 and RCM_triCNN on combining set1)

### Model performance evaluation on the testing set

To ensure an unbiased evaluation of the model performance, a testing set containing 2,226 LS and BS exon pairs (**see Fig. 2a**) was utilized to assess model performance. ROC curves were plotted to visualize the performance of the retrained optimal base, RCM_triCNN, and combined models on the testing set, as shown in **Fig. 8**. Specifically, for each training set (where the base model and RCM_triCNN model were trained on) and each corresponding combining set (where the combined model was trained on), the ROC curve based on the model performance on the testing set was depicted. In total, 15 models were compared, including 3 optimal base model1s, 3 optimal base model2s, 3 optimal RCM_triCNN models trained on each of the 3 training sets, 3 optimal combined models of base model1 with RCM_triCNN, and 3 optimal combined models of base model2 with RCM_triCNN trained on each of the 3 corresponding combining sets. In theory, a perfect classifier would produce an AUC (area under the curve) value of 1, and a random classifier would produce an AUC value of 0.5, and the closer the AUC value to 1 the better the model performance. As shown in **Fig. 8**, the AUC value for all base models exceeded 0.91, indicating strong model performance. While the AUC values for RCM_triCNN models were less than 0.8, integration with base models improved the combined model performance, as measured by additional performance metrics shown in **Table 1.** For instance, the best ACC of 0.8813 was achieved by the combined base model2, where integration with RCM_triCNN increased the ACC of the base model2 from 0.8750 to 0.8813. Importantly, integration with RCM_triCNN improved the specificity of both base models (*e.g.*, from 0.9316 to 0.9658 for base model1, and from 0.8974 to 0.9451 for base model2 on combining set2).

**Fig. 8.**
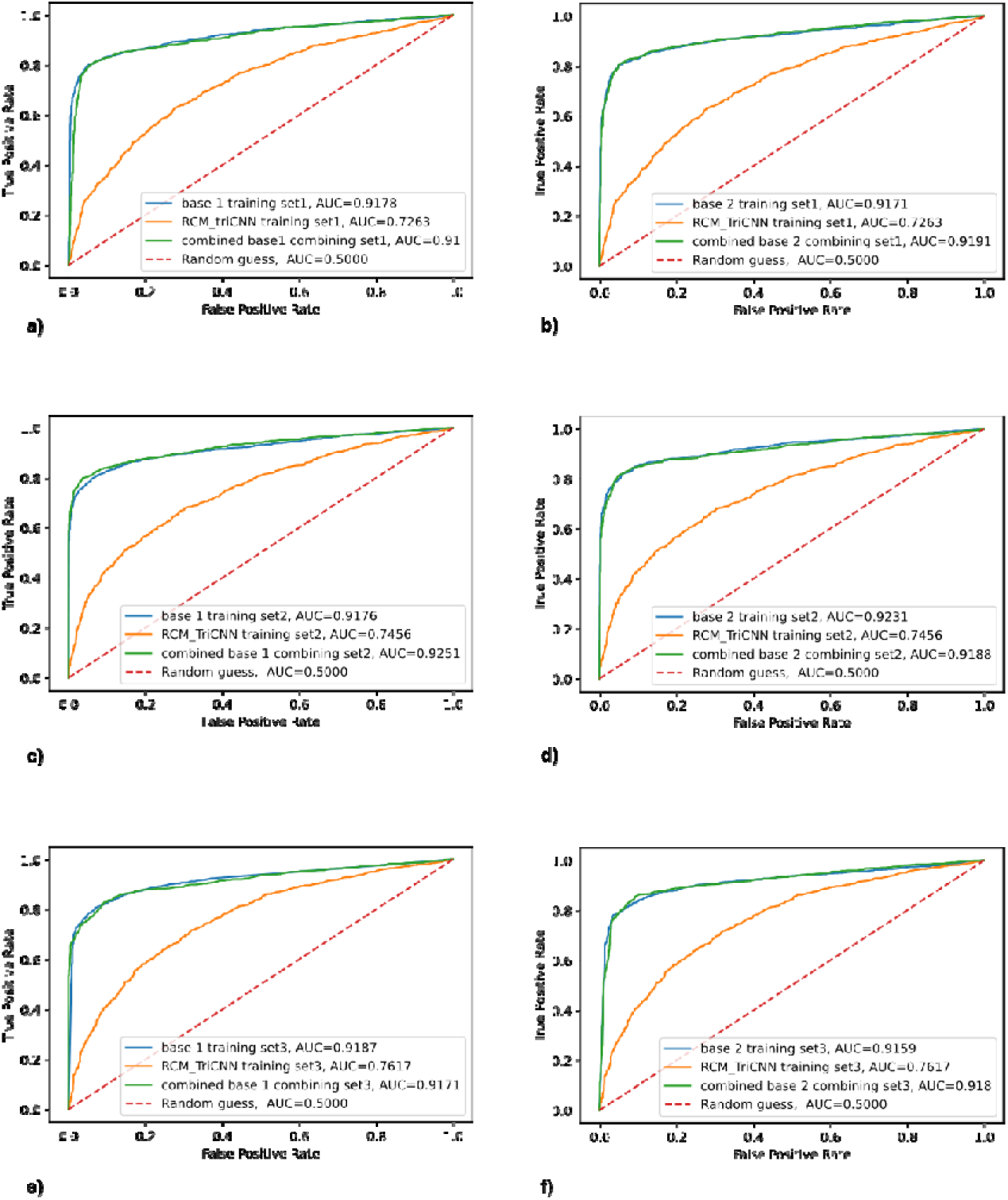
ROC Curve for the retrained optimal base, RCM_triCNN, and combined models on the testing set. a): ROC curves for the base model1 (trained on the training set1), RCM_triCNN model (trained on the training set1) and combined base model1 (same model weights as base model1 and RCM_triCNN with additional layers trained on combining set1) on the test dataset. b): ROC curves for the base model2 (trained on the training set1), RCM_triCNN model (trained on the training set1) and combined base model2 (same model weights as base model2 and RCM_triCNN with additional layers trained on combining set1) on the test dataset. c): ROC curves for the base model1 (trained on the training set2), RCM_triCNN model (trained on the training set2) and combined base model1 (same model weights as base model1 and RCM_triCNN with additional layers trained on combining set2) on the test dataset. d): ROC curves for the base model2 (trained on the training set2), RCM_triCNN model (trained on the training set2) and combined base model2 (same model weights as base model2 and RCM_triCNN with additional layers trained on combining set2) on the test dataset. e): ROC curves for the base model1 (trained on the training set3), RCM_triCNN model (trained on the training set3) and combined base model1 (same model weights as base model1 and RCM_triCNN with additional layers trained on combining set3) on the test dataset. f): ROC curves for the base model2 (trained on the training set3), RCM_triCNN model (trained on the training set3) and combined base model2 (same model weights as base model2 and RCM_triCNN with additional layers trained on combining set3) on the test dataset.

## Discussion

Despite the recognized functional importance of circRNAs in various biological processes, the molecular mechanisms governing their biogenesis remain incompletely understood. Among the proposed mechanisms, back-splicing (BS) is considered a prominent pathway for exonic circRNA formation^3^. Both cis-elements and trans-factors are important in this process. For example, experimental evidence has underscored the crucial role of reverse complementary matches (RCMs) in BS-mediated circRNA formation^34^. Proteins such as Quaking (QKI) and MBL/MBNL1 (muscleblind) have been identified as splicing factors that can promote circRNA formation by facilitating proximity between splice sites^6^. Several other RNA-binding factors including TNRC6A, NF90/NF110, ESRP1, FUS, HNRNPL, and DHX9 have also been implicated in circRNA formation^35,36^, although their roles may vary depending on the context. Despite these advancements, the underlying mechanism governing the circRNA formation is still elusive.

While machine learning approaches have been successfully applied to classify circRNAs from lncRNAs with increasing prediction accuracy^12^, efforts to elucidate the formation of exonic circRNAs using machine learning techniques have been limited due to the lack of properly labeled, high-quality datasets. Therefore, in this study, we employed a comprehensive data integration approach to curate a dataset of BS and LS exon pairs with evidence-based filtering. Our results demonstrated that most of the BS exon pairs from the current circRNA database showed evidence of LS (*e.g*., circRNA exon pairs reduced significantly from 386,363 to 15,030 upon filtering with the LS exon pairs dataset), highlighting the importance of stringent data selection criteria for applications of machine learning approaches to understand circRNA biogenesis.

Based on our high-quality datasets, we designed the CircCNNs framework and explored the weights of trained models to better understand the potential molecular mechanisms of exonic circRNA formation. Different from the previous study^19^, we designed a CNN framework that includes two sets of CNN structures for both upstream and downstream junction sequences of BS and LS exon pairs. This approach aims to discover potentially distinctive sequence motifs for RNA binding proteins (RBPs) involved in circRNA formation. As a comparison, we also trained the model (base model1) with a similar structure as the previous study^19^. In addition to modeling the junction sequences, we also created models that only take in RCM information of exon pairs. To determine the RCMs crossing and within the flanking intron regions of splicing sites (exon pairs), we created and implemented an efficient novo numerical approach to mitigate the computational cost for large kmer length. For comprehensive RCM determinations, 3 separate flanking window sets (ranging from 100 to 3,000 bps) of splicing sites were used. To retain the spatial information of RCMs, RCM frequency was converted into corresponding RCM kmer pair distribution matrix based on segmented flanking regions of equal distance. With the RCM data, two separate RCM models were created to investigate the relative importance between the RCMs crossing and within flanking regions of the splicing sites (see **Fig. 3** for RCM determination, **Fig. 4a** and **4b** for RCM_crossCNN vs RCM_triCNN structures respectively) for model performance. Recognizing the complementary information contents provided by both junction sequences and RCMs, we integrated these two models into combined models to further improve the model performance. To investigate the effects of training sets on the model performance as well as on the motif discovery, we created 3 different training sets with incrementally larger sizes as well as 3 corresponding combining sets for model integrations (see **Fig. 2a**).

CNNs have been extensively used in genomic sequence modeling, owing in part to their capability to extract potentially important motifs from trained models^37^. In this study, we followed the motif visualization technique^33^ outlined in a previous study to uncover genes potentially pivotal for BS or LS. Despite different model structures trained on slightly different datasets, genes, such as RBFOX1, ZC3H10, ENOX1, and MBNL1, emerged as significant for both LS and BS across all base models (highlighted with a red asterisk in **Fig. 6a**). Some of these genes play key roles as RNA splicing regulators and may be implicated in BS or LS processes. For instance, RBFOX1 (RNA Binding Fox-1 Homolog1) protein is involved in alternative splicing by binding to 5’-UGCAUGU-3’ elements^38^. MBNL1 (Muscleblind Like Splicing Regulator 1) encodes an alternative splicing regulator that can bind to stem-loop structure within the polypyrimidine tract of TNNT2 intron to inhibit spliceosome complex A formation^38^; notably, it has also been implicated in promoting circRNA formation^6^. For genes potentially exclusive for BS, base model1 identified HuR, RBMS1, HNRPLL, RBM42, QKI and PTBP1, while upstream-CNN of base model2 identified RBM24, LIN28A, SRSF2, and SRSF7; Downstream-CNN of base model2 identified PABPN1, RBM28, ZNF638, MATR3, ESRP2 and ANKHD1. Among these genes, Quaking (QKI) has been shown to promote circRNA formation^6^, while ESRP1 (epithelial splicing regulatory protein 1) has been implicated in circRNA formations. Notably, QKI was identified as important for both LS and BS by upstream- and downstream-CNN of base model2 (see **Fig. 6a**). Given the rigorous criterion employed in curating the BS and LS datasets, the experimentally verified BS-associated genes collectively identified in our study underscore the high quality of our curated datasets and the utility of our model structures in elucidating exonic circRNA formations.

On average, the base model2 demonstrated slightly superior performance compared to the base model1 in all 3 different training sets. Regarding the RCM models, RCM_triCNN consistently outperformed RCM_crossCNN (see **Fig. 5b-d)**. However, their performance remained notably lower than that of the base models (see **Fig. 5**, **Fig. 8, and Table 1**), suggesting the potential dominance of trans-factors, such as RBPs, over cis-elements, represented by RCMs in the circRNA formation. This observation aligns with the previous studies that while RCMs can enhance exonic circRNA formation, they are not essential^39^. Past studies on distinguishing between lncRNAs and circRNAs underscored the significance of RCMs crossing the flanking intron regions of the splicing sites as a key discriminatory feature^11^. Here, we introduce a novel approach by incorporating RCMs from each flanking region of the splicing sites, which notably enhances model performance in BS and LS exon pair classification. Interestingly, varying window sizes in the RCM models had only marginal effects on model performance, underscoring the importance of RCM information proximal to the splicing sites in BS.

**Table 1:** Summary of model performance on the testing set.

|  | ACC | Specificity | Recall | Precision | FPR | FNR | F1 | MCC | AUROC | AUPRC |
| --- | --- | --- | --- | --- | --- | --- | --- | --- | --- | --- |
| base 1 training set1 | 0.8718 | 0.9406 | 0.8027 | 0.9307 | 0.0594 | 0.1973 | 0.8620 | 0.7507 | 0.9178 | 0.9362 |
| RCM_triCNN training set1 | 0.6697 | 0.6535 | 0.6860 | 0.6632 | 0.3465 | 0.3140 | 0.6744 | 0.3396 | 0.7263 | 0.7265 |
| combined base 1 combining set1 | 0.8709 | 0.9442 | 0.7973 | 0.9343 | 0.0558 | 0.2027 | 0.8604 | 0.7498 | 0.91 | 0.9205 |
| base 2 training set1 | 0.8714 | 0.9541 | 0.7882 | 0.9447 | 0.0459 | 0.2118 | 0.8594 | 0.7530 | 0.9171 | 0.9381 |
| combined base 2 combining set1 | 0.8764 | 0.9505 | 0.8018 | 0.9416 | 0.0495 | 0.1982 | 0.8661 | 0.7610 | 0.9191 | 0.9377 |
| base 1 training set2 | 0.8678 | 0.9316 | 0.8036 | 0.9212 | 0.0684 | 0.1964 | 0.8584 | 0.7415 | 0.9176 | 0.9398 |
| RCM_triCNN training set2 | 0.6873 | 0.6949 | 0.6796 | 0.6890 | 0.3051 | 0.3204 | 0.6843 | 0.3746 | 0.7456 | 0.7565 |
| combined base 1 combining set2 | 0.8800 | **0.9658** | 0.7937 | **0.9585** | **0.0342** | 0.2063 | 0.8683 | **0.7712** | **0.9251** | **0.9462** |
| base 2 training set2 | 0.8750 | 0.8974 | **0.8525** | 0.8920 | 0.1026 | **0.1475** | 0.8718 | 0.7507 | 0.9231 | 0.9436 |
| combined base 2 combining set2 | **0.8813** | 0.9451 | 0.8172 | 0.9367 | 0.0549 | 0.1828 | **0.8729** | 0.7688 | 0.9188 | 0.9403 |
| base 1 training set3 | 0.8673 | 0.9253 | 0.8090 | 0.9150 | 0.0747 | 0.1910 | 0.8588 | 0.7395 | 0.9187 | 0.9269 |
| RCM_triCNN training set3 | 0.6958 | 0.6967 | 0.6950 | 0.6950 | 0.3033 | 0.3050 | 0.6950 | 0.3917 | 0.7617 | 0.7564 |
| combined base 1 combining set3 | 0.8615 | 0.9154 | 0.8072 | 0.9047 | 0.0846 | 0.1928 | 0.8532 | 0.7270 | 0.9171 | 0.9395 |
| base 2 training set3 | 0.8709 | 0.9289 | 0.8127 | 0.9191 | 0.0711 | 0.1873 | 0.8626 | 0.7468 | 0.9159 | 0.9287 |
| combined base 2 combining set3 | 0.8745 | 0.9334 | 0.8154 | 0.9241 | 0.0666 | 0.1846 | 0.8663 | 0.7542 | 0.9180 | 0.9295 |

Leveraging potentially complementary information from nearby junction sequences and RCMs, the integration of the base models with RCM_triCNN models across different combining sets enhanced model performance as shown in **Fig. 7**. With a spectrum of metrics including ACC, MCC, and AUC values, all base models showed good performance, with further enhancement observed upon integrating the RCM_triCNN model (see **Table 1**). DeepCirCode^19^ achieved a sensitivity of 0.9214, with a specificity of 0.7684 in their proposed dataset, implying the potentially high false positive predictions. In this study, with our curated dataset, the lowest and highest specificity achieved among all base models were 0.8974 and 0.9541, respectively. We also found that integration of RCM_triCNN increased the specificity of the base models significantly, particularly prominent on the combining set2 (see **Table 1**). Overall, integration with RCM_triCNN led to increased precision and reduced FPR for both base models, with a slight tradeoff for model sensitivity. The CircCNNs framework introduced here holds the potential for mitigating high false positives in both existing circRNA detection pipelines processing RNA-seq data and circRNA databases.

## Acknowledgments

We are grateful for the support from Dr. Jens Mueller with the help of using the Miami Redhawk cluster for model training.

## Author contributions

CW designed the study, implemented the model, analyzed the data, and wrote the manuscript; CL provided valuable feedback and helped with the clarification of the entire manuscript.

## Data availability statement

The data supporting the findings of this study are available within the article [and/or] its supplementary materials.

