## Supplemental figure1 for "CircCNNs, a convolutional neural network framework to better understand the biogenesis of exonic circRNAs"

**Department of Biology, Miami University, Oxford, Ohio 45056**

* Corresponding author

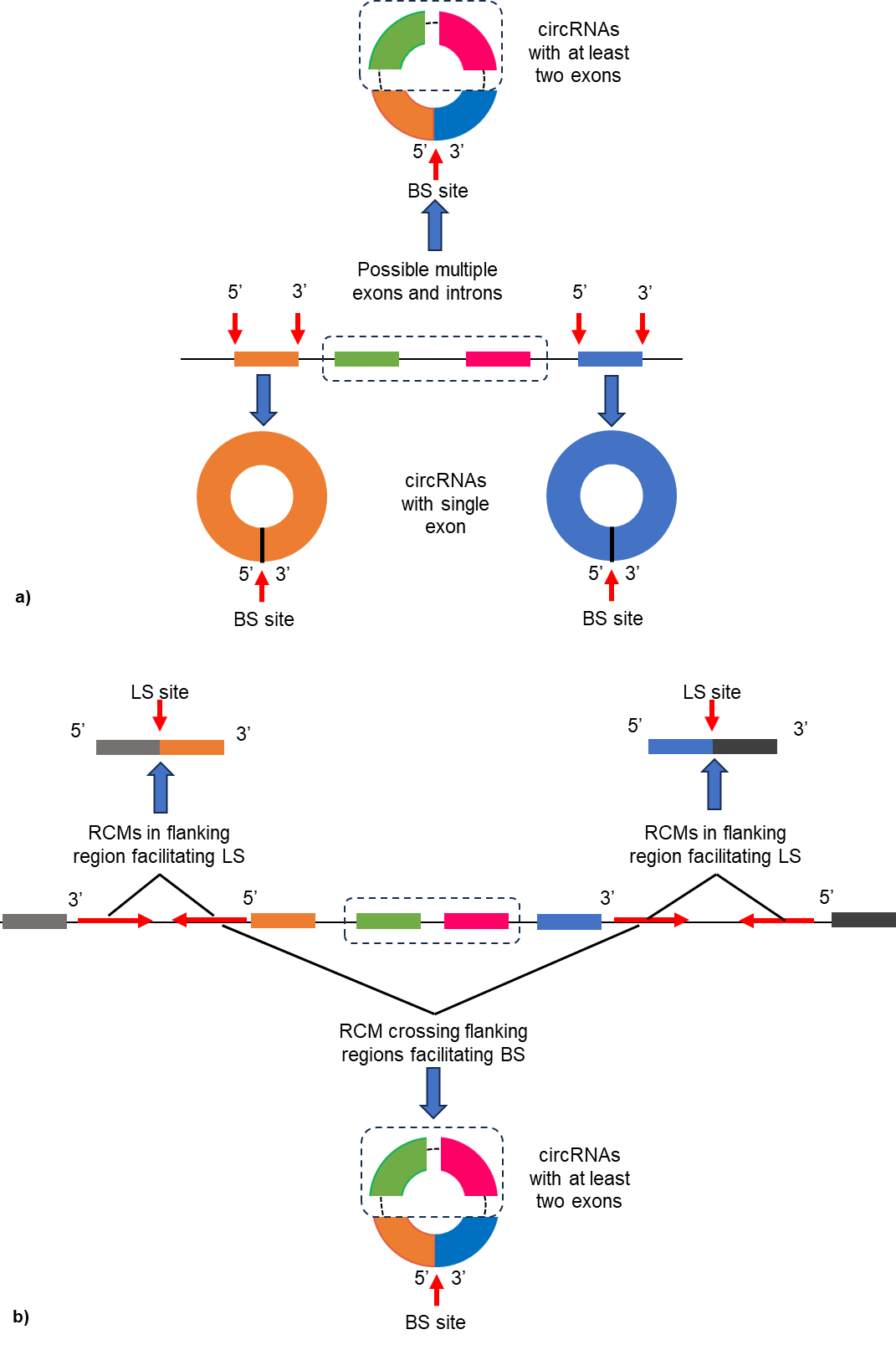

**Fig. S1** a): possible circRNAs formed by BS; b): competition of RCM crossing and within flanking intron regions of the exon pair could potentially influence the outcome of BS or LS
